# Olfactory bulb stimulation mitigates Alzheimer’s-like disease progression

**DOI:** 10.1101/2024.03.03.583116

**Authors:** Morteza Salimi, Milad Nazari, Payam Shahsavar, Samaneh Dehghan, Mohammad Javan, Javad Mirnajafi-Zadeh, Mohammad Reza Raoufy

## Abstract

**Background:** Deep brain stimulation (DBS) has demonstrated potential in mitigating Alzheimer’s disease (AD). However, the invasive nature of DBS presents challenges for its application. The olfactory bulb (OB), showing early AD-related changes and extensive connections with memory regions, offers an attractive entry point for intervention, potentially restoring normal activity in deteriorating memory circuits.

**Aims:** Our study examined the impact of electrically stimulating the OB on working memory as well as pathological and electrophysiological alterations in the OB, medial prefrontal cortex, hippocampus, and entorhinal cortex in amyloid beta (Aβ) AD model rats.

**Methods:** Male Wistar rats underwent surgery for electrode implantation in brain regions, inducing Alzheimer’s-like disease. Bilateral olfactory bulb (OB) electrical stimulation was performed for 1 hour daily to the OB of stimulation group animals for 18 consecutive days, followed by evaluations of histological, behavioral, and local field potential signal processing.

**Results:** OB stimulation counteracted Aβ plaque accumulation and prevented AD-induced working memory impairments. Furthermore, it prompted an increase in power across diverse frequency bands and enhanced functional connectivity, particularly in the gamma band, within the investigated regions during a working memory task.

**Conclusion:** This preclinical investigation highlights the potential of olfactory pathway-based brain stimulation to modulate the activity of deep-seated memory networks for AD treatment. Importantly, the accessibility of this pathway via the nasal cavity lays the groundwork for the development of minimally invasive approaches targeting the olfactory pathway for brain modulation.

## 1 Introduction

The escalating prevalence of Alzheimer’s disease (AD) as an age-related ailment poses substantial global challenges.^1^ AD diminishes individuals’ and their families’ quality of life and strains healthcare systems.^2^ Thus, early-stage diagnostic and intervention methods are crucial for alleviating the social and financial burdens linked with AD.^3^ Mild Cognitive Impairment (MCI), denoting a cognitive decline without a significant impact on daily activities.^4^ It’s imperative to explore ways to prevent the progression from MCI to AD, a transition occurring in roughly 80% of MCI cases over about 6 years.^5^

AD, characterized by Aβ accumulation in different brain regions^6^ disrupts brain homeostasis, resulting in diverse cognitive impairments, notably memory disorders.^7^ Pathophysiological changes like neurofibrillary tangles and neuritic plaques initially manifest in regions including the olfactory bulb (OB), entorhinal cortex (EC), and hippocampus (HPC).^6^ An early symptom of AD is a diminished or altered sense of smell preceding other cognitive impairments.^8,9^ The OB is intricately connected to limbic regions and plays a role in memory control within a network involved memory-related functions.^10-12^ The EC acts as the gateway for information entering and exiting the hippocampus.^13^ It also serves as an interface between the medial prefrontal cortex (mPFC) and HPC for the rapid encoding of new memories.^14^ Communication between the PFC and HPC is crucial for the encoding, processing, and execution of memory.^15^ mPFC exerts control over HPC-dependent memories and providing hippocampal top-down modulation.^15^ OB plays a coordinating role in the cross-talk in this circuit particularly during working memory tasks.^11^ Disruptions in the OB-EC-HPC-mPFC network, such as plaque accumulation and changes in cell viability during MCI, can impair memory performance.^16-18^ Considering early AD biomarker presence in the OB and its vital role in this network, targeting the OB holds promise for preventive interventions.

The efficacy of deep brain stimulation (DBS) in slowing AD progression has been investigated in both animal and human studies.^19^ High-frequency DBS in specific brain regions improves cognitive functions and reduces AD pathology in animal model.^20-23^ In clinical trials, DBS in the fornix and nucleus basalis of Meynert has shown significant efficacy.^19,24^ DBS could enhance memory by reducing pathological hallmarks like beta-amyloid, increasing cell survival, regulating neural networks, and promoting neurogenesis.^19^ However, a standardized stimulation protocol specifying the optimal brain area is still lacking. Given its intricate connections with various memory-associated regions, the OB emerges as a compelling candidate for targeted brain stimulation, especially in the early MCI stage when electrophysiological and pathophysiological hallmarks begin to appear. Our previous studies demonstrated that OB stimulation not only improved impaired working memory caused by mechanical ventilation but also acted preventively against HPC inflammation and apoptosis.^25,26^ Other studies have shown the effectiveness of OB stimulation using odorants in enhancing cognitive functions in AD patients.^27^ Overall, we propose that OB stimulation in the early stages of AD can prevent memory disorders, amyloid plaque accumulation, and deterioration of memory-related networks. In this study, we aimed to electrically stimulate the OB while inducing an AD model in rats and examined the stimulation’s impact on working memory performance and potential pathophysiological alterations related to AD progression within the OB-EC-HPC-mPFC circuit.

## 2 Materials and methods

### 2.1 Animals

The experiments were performed on pathogen-free male Wistar rats. The rats weighed 210-230 g. All animals were obtained from Tarbiat Modares University (Tehran, Iran) and kept at 21 ± 2 °C on a 12-hour light-dark cycle. They had unrestricted access to food and water. The Ethics Committee of the Faculty of Medical Sciences, Tarbiat Modares University approved all experimental procedures (IR.MODARES.REC.1398.037).

### 2.2 Surgery and Electrode implantation

Electrode implantation and i.c.v. injection were carried out in accordance with our previous protocol.^28^ Briefly, after anesthetizing, the animals were placed on the stereotaxic apparatus, and then the skull was exposed for finding the coordinates in the Paxinos atlas. Based on the reference points in the Paxinos atlas, we found the coordinates for OB (AP: 8.5 mm, ML: ± 1 mm, DV: − 1.5 mm), i.c.v. (AP: 0.8 mm, ML: ± 1.6 mm, DV: − 3 mm), mPFC (AP + 3.2 mm, ML −0.6 mm, DV −3.6 mm), dHPC (AP: − 3.6 mm, ML: − 2.2 mm, DV: − 2.7 mm), vHPC (AP:−4.92 mm, ML −5.5 mm, DV −7.5 mm) and EC (AP: − 7.04 mm, L: − 5.5 mm, DV: − 6.5 mm). We drilled holes in the skull and stainless-steel twisted recording electrodes (127 μm in diameter, A.M. System Inc., USA) were implanted in the five regions of interest. To confirm that electrodes were located correctly, we removed the rats’ brains after perfusion and then kept a hemisphere in 4% paraformaldehyde for 48 h. Finally, a 200 μm coronal section was taken to visually compare with matching slices in the Paxinos and Watson rat brain atlas^29^ (Fig. 1).

**Figure 1.**
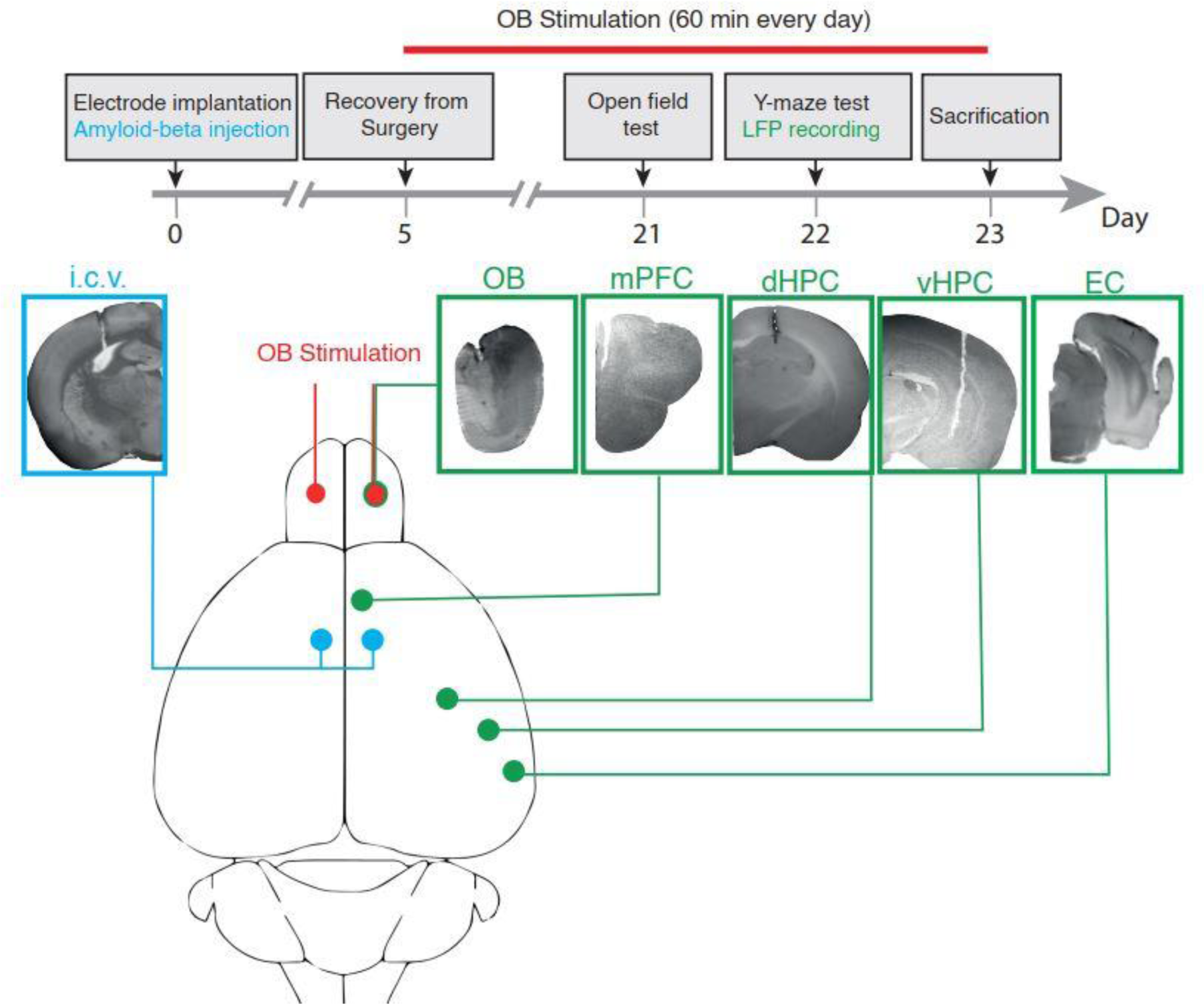
Protocol of the study. (Upper panel) indicates bilateral Aβ or saline injections and unilateral electrode implantations were carried out on day zero. After 5 days of recovery, electrical stimulation bilaterally delivers for 1 hour daily to the OB of stimulation group animals for 18 consecutive days. On days 21 and 22, the open field test and Y-maze tasks were performed, respectively. (Lower panel) displays schematic and histological verification of i.c.v. injection as well as electrode implantation. OB, olfactory bulb; mPFC, medial prefrontal cortex; dHPC, dorsal hippocampus; vHPC, ventral hippocampus; EC, entorhinal cortex.

### 2.3 AD model induction

In order to induce Alzheimer’s-like disease, we used Aβ1–42 (Cat No. A9810, Sigma-Aldrich, USA) dissolved in normal saline at a concentration of 4 μg/μl. After allowing the solutions to sit at room temperature for 3 days, we bilaterally injected either 2 μl of saline or 2 μl of Aβ1–42. The injections were administered continuously over 5 minutes using a microinfusion pump (Stoelting, Lane Dale, IL, USA) connected to 25-gauge stainless steel cannulas. Therefore, four groups of animals were included in the current study (Saline group: animals which received i.c.v. saline; Saline+Stim: animals which received i.c.v. saline accompanied by OB stimulation; Aβ: animals which received i.c.v. Aβ; Aβ+Stim: animals which received i.c.v. Aβ accompanied by OB stimulation).

### 2.4 Electrical Stimulation and electrophysiological recording

Five days after surgery, the animals underwent bilateral OB electrical stimulation. The stimulation was delivered continuously for 1 hour daily (130 Hz, 90 μs pulse width, 50 μA amplitude). Electrical artifacts were monitored to confirm stimulation delivery throughout the 1-hour sessions (Fig. 2). For both stimulation and local field potentials (LFP) recording, we applied 127 μm diameter electrodes (A.M. System Inc., USA) connected to a miniature buffer head stage having high-input impedance (BIODAC-A, TRITA Health Technology Co., Tehran, Iran). The obtained signals were amplified with a 1000x gain, low-pass filtered < 250 Hz, 50 Hz notch filter, and digitized at 1 kHz through a controller (BIODAC-ESR18622, TRITA Health Technology Co., Tehran, Iran).

**Figure 2.**
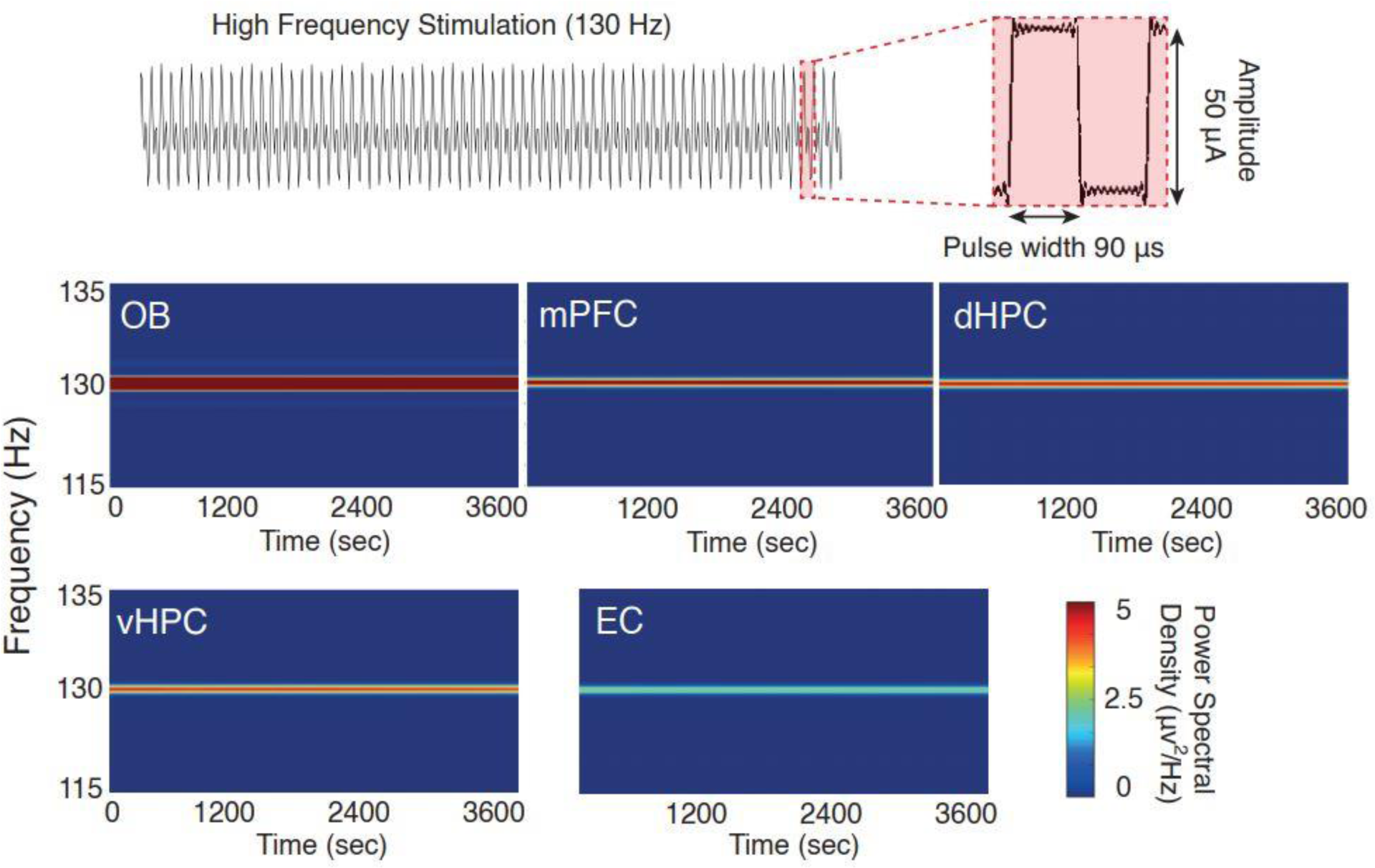
OB electrical stimulation. (Upper panel) A continuous 130 Hz electrical pulse with 90 μs pulse width and 50 μA amplitude was delivered bilaterally into the OBs. (Lower panel) Artifact signals in five brain regions during 1 hour at the same frequency of OB electrical stimulation. OB, olfactory bulb; mPFC, medial prefrontal cortex; dHPC, dorsal hippocampus; vHPC, ventral hippocampus; EC, entorhinal cortex.

### 2.5 Histological Assessments

Immunofluorescence was conducted as follows: rats were subjected to transcranial perfusion using cold phosphate-buffered saline (PBS) with a concentration of 10 mM and a pH of 7.4. This was followed by perfusion with a solution containing 4% paraformaldehyde in PBS, while the rats were under urethane anesthesia at a dose of 1.2 g/kg. After perfusion, the brains were carefully removed and post-fixed in a 4% paraformaldehyde-PBS solution. Subsequently, the brain samples were cryopreserved by immersing them in a 30% sucrose solution prepared in PBS for a period of three days at a temperature of -4 °C. Following cryopreservation, the brain samples were embedded in an optimum cutting temperature (OCT) compound (Sakura Finetek, USA), frozen, and then stored at -80 °C. Coronal sections measuring 8 μm in thickness were obtained from OB, EC, PFC, vHPC, and dHPC. These sections were prepared using a freezing microtome (Histo-Line Laboratories, Italy) and mounted onto superfrost plus slides.

For immunofluorescent labeling, frozen slides were allowed to air dry at room temperature (RT) for 30 minutes. Subsequently, they were rinsed with a solution of 0.05% Tween-80 in PBS for 15 minutes and then permeabilized using a 0.2% Triton X100 solution (Sigma-Aldrich, USA) in PBS. To prevent non-specific binding, the slides were treated with a blocking solution containing PBS, 0.2% Triton-X100, and 10% normal serum for one hour at RT. The primary antibodies, specifically anti-Aβ antibodies (A8717, diluted at 1:700), were applied in the blocking solution and left to incubate overnight at 4 °C. On the following day, after thorough rinsing with PBS, the slides were incubated with appropriate fluorescent-labeled secondary antibodies (goat anti-rabbit, Alexa Fluor® 488, diluted at 1:2000, ab150077) for one hour at RT. To visualize cell nuclei, 4’,6-diamidino-2-phenylindole (DAPI; Invitrogen Corp., USA) was used for counterstaining. The slides were then mounted, and images were captured using Olympus fluorescence microscopy equipped with a DP72 camera (BX51 TRF, USA).

The quantification of Aβ immunoreactivity involved measuring fluorescence intensity, which was subsequently normalized using the mean value from the Saline group. Data collected from individual animals within each group was first averaged based on three sections, and then further averaged for each of the three animals in the treatment group. The assessment of Aβ fluorescence intensity was conducted by calculating the mean gray value in specific regions of interest, including the OB, EC, PFC, vHPC, and dHPC, using Fiji software. Each obtained value was then normalized to the mean intensity of the tissue background and expressed as a percentage relative to the Saline group.

Nissl staining was employed to assess the survival of cells within the OB across all experimental groups. The procedure adhered to the manufacturer’s guidelines and involved the following steps: sections were rehydrated through a series of graded alcohol solutions (96%, 80%, and 70%), followed by staining with a 0.1% Cresyl Fast Violet solution (Merck, Germany) at RT for 5 minutes. Subsequently, the sections were rinsed, dehydrated via a graded alcohol series (70%, 80%, 96%, and 100%), cleared using xylene, cover-slipped using Entellan (Merck, chemical, Germany), and then photographed. Images were captured from non-overlapping, consecutive microscopic fields using an Olympus BX-51 microscope equipped with a DP72 camera at 400x magnification. To assess cell count, a random grid measuring 50 μm x 50 μm was applied to the images and the count was conducted across seven squares. Fiji software was utilized for cell number analysis.

### 2.6 Signal processing

We selected the signals of LFP during spontaneous alterations when animals explored the center of the maze (Red triangle in Fig. 5). These time periods were selected for further analyses. Finally, the signal processing assessment during correct trials was averaged per animal and compared with wrong trials within the group and between groups. LFPs were simultaneously obtained from five regions of interest. Signals were preprocessed using the EEGLAB toolbox for noise rejection and baseline correction.^12^ Power spectral density (PSD) was computed using the Welch’s periodogram function in MATLAB.^30^ To plot connection edges within five brain regions, we first calculated the coherence using mscohere function in MATLAB, and then any pairwise regions that passed the designated threshold of coherence were connected. The color of connected lines represents the value of coherence between brain regions.

### 2.7 Statistical Analysis

GraphPad Prism (version 6.0) was applied for statistical analyses and for creating graphs. We examined the normality distribution of the data via the Kolmogorov–Smirnov test. A two-way ANOVA test was used to compare significant changes within and between groups. For all comparisons, post hoc Bonferroni correction was applied to determine specific group differences. The p-value less than 0.05 was considered statistically significant.

## 3 Results

### 3.1 OB stimulation prevents working memory impairment in AD-like animals

The spontaneous alternation Y-maze is a widely employed method for assessing working memory in rodents.^31^ Rodents naturally tend to explore novel environments. In this task, rats sequentially explore three arms, necessitating them to recall the most recently visited arm and then select a new one.^31^ Animals with impaired working memory are likely to show reduced spontaneous alternation since they are not able to remember which arm was visited last.^32^ Working memory impairment is frequently observed in patients with AD.^33^ Our study aimed to investigate the induction of an Alzheimer’s-like condition using Aβ i.c.v. injection, which could potentially impact working memory. We found Aβ animals exhibited significantly reduced working memory compared to controls (Aβ vs. Saline, p < 0.001). Interestingly, OB stimulation for one hour a day prevented working memory impairment in Aβ animals (Aβ+Stim vs. Aβ, p < 0.001). However, OB stimulation in the control animals did not produce a significant effect on working memory when compared to the animals that didn’t receive stimulation (Fig. 3). No significant alteration was found in locomotor activity among the four experimental groups. Therefore, our findings suggest that OB stimulation can mitigate working memory impairment in AD-like animals.

**Figure 3.**
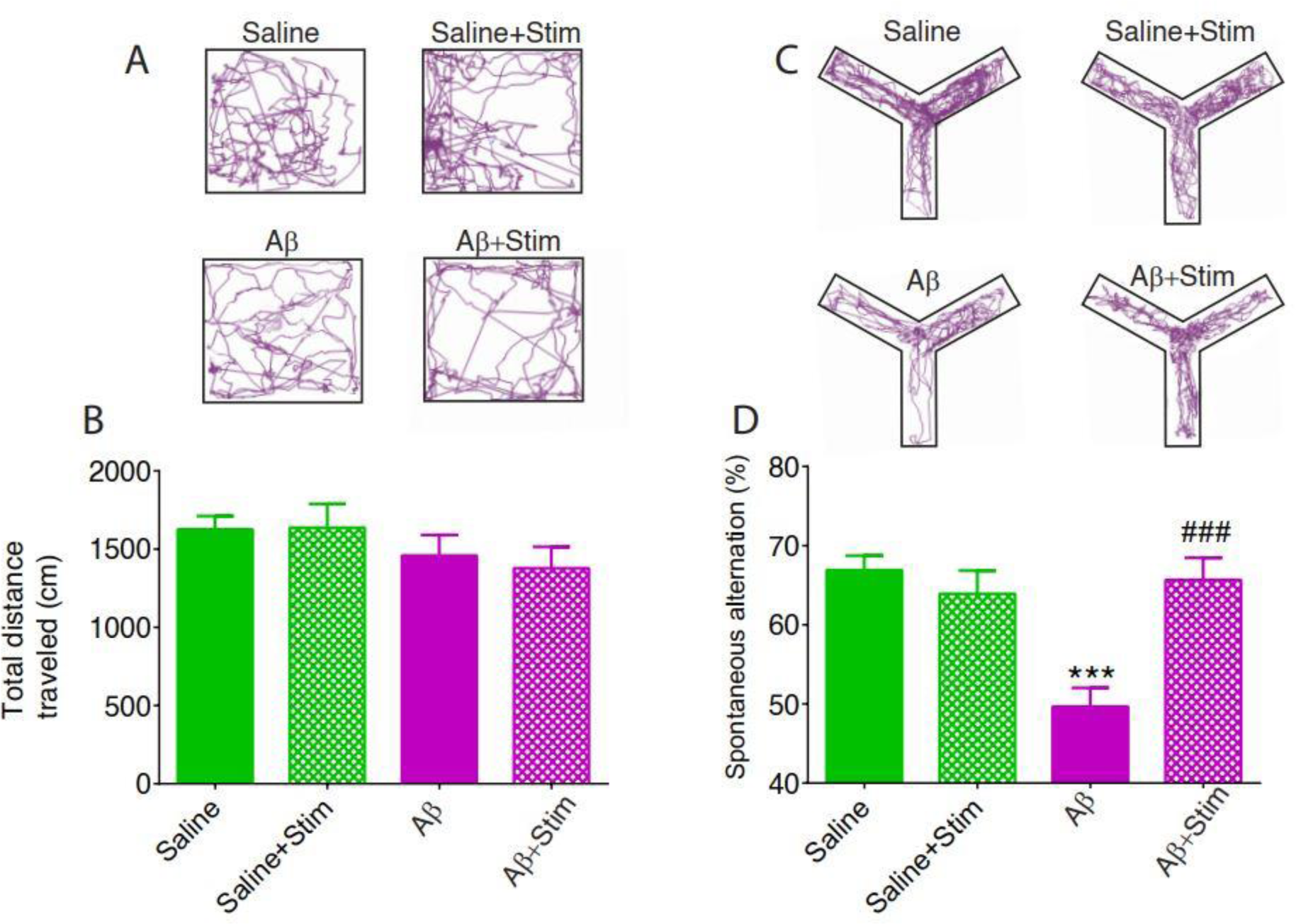
Locomotor activity and working memory task. (A, C) representative example of locomotor activity in the open field and spontaneous alteration in Y-maze. (B) No significant changes were found in the traveled distance as an indicator of locomotor activity. (D) Aβ significantly decreased the correct arm visits, which indicates a decline in working memory performance compared to the Saline group. OB stimulation prevented working memory impairment in Aβ animals. n=6; Data were analyzed by two-way ANOVA; *** p < 0.001 compared to Saline or Saline+Stim; ### p < 0.001 compared to Aβ.

### 3.2 OB stimulation mitigates the cellular loss in the OB and the Aβ aggregation in the brain regions of AD-like animals

In this study, we examined the impact of Aβ injection and OB stimulation on cellular survival of OB and plaque accumulation in various brain regions (Fig. 4). Injection of Aβ i.c.v. route led to a significant reduction in the number of surviving cells within the OB (Aβ vs. Saline, p < 0.01), underscoring the detrimental effects of Aβ on cellular viability. However, when considering OB stimulation, a noteworthy contrast emerged. Aβ animals subjected to OB stimulation displayed a significant reduction in cell loss (Aβ+Stim vs. Aβ, p < 0.05). This finding suggests a potential protective effect of OB stimulation on Aβ-induced cellular loss.

**Figure 4.**
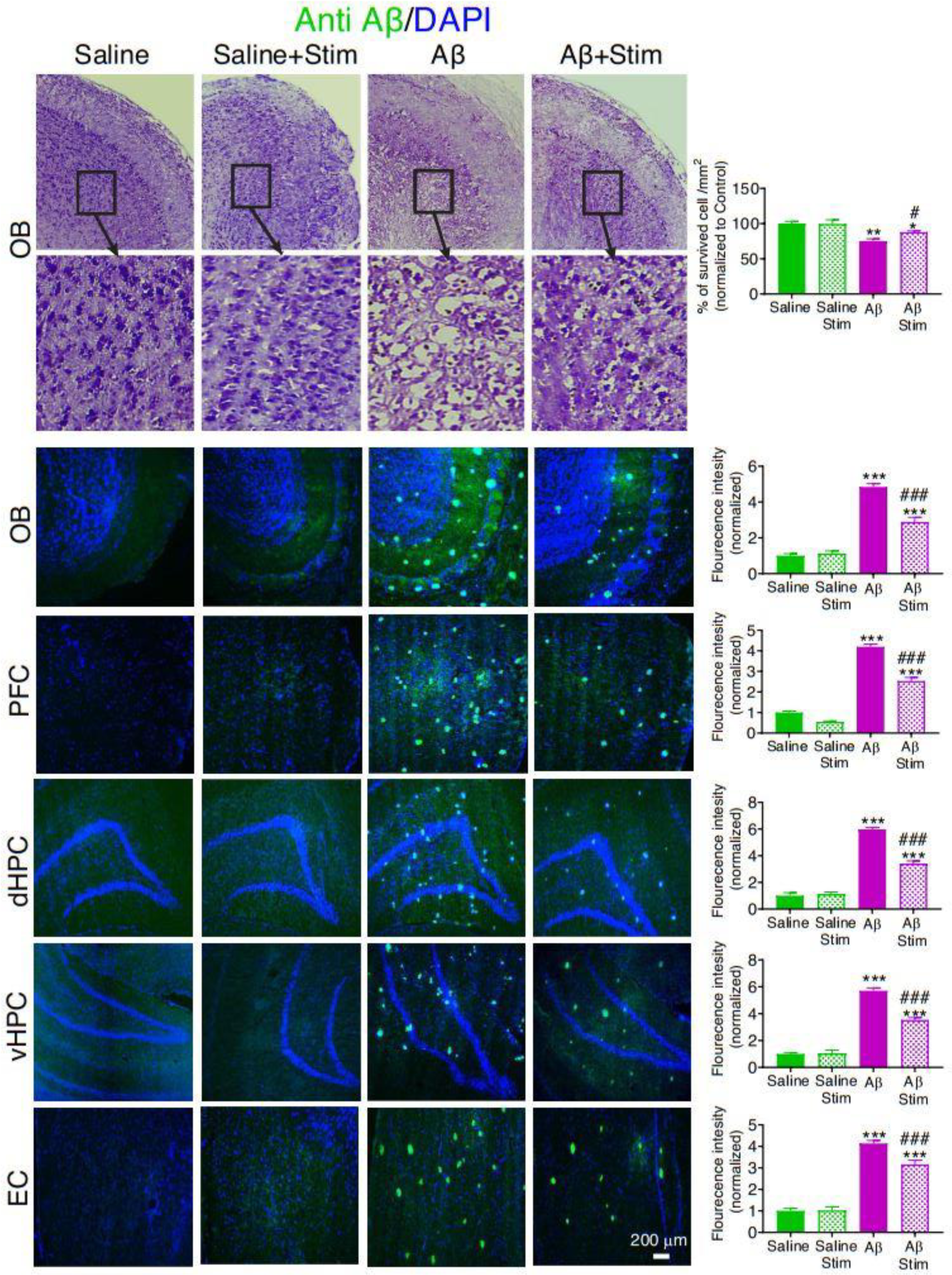
Histological assessments. Representative sample of survived cells in the OB (top) and sample of immunostaining sections groups of study (bottom). Injection of Aβ via the i.c.v. led to a significant decrease in the number of surviving cells within the OB. Interestingly, stimulation of the OB resulted in a notable reduction in cell loss in Aβ animals. Immunostaining using anti-Aβ1-42 antibodies revealed a substantial accumulation of plaques in animals exposed to Aβ. Remarkably, OB stimulation led to a reduction in plaque accumulation in Aβ animals not only in the OB but also in the PFC, dHPC, vHPC, and EC. n=6; Data were analyzed by two-way ANOVA; *p < 0.05, **p < 0.01,***p < 0.001 compared to Saline or Saline+Stim; #p < 0.05, ###p < 0.001 compared to Aβ group.

Immunostaining targeting anti-Aβ1-42 was employed to probe plaque accumulation in the brain. As shown in figure 4, The results revealed a substantial increase in plaque accumulation within Aβ animals when compared to the saline groups (Aβ vs. Saline, p < 0.001). This heightened plaque accumulation underscores the pathological impact of Aβ, potentially contributing to cognitive impairment. Intriguingly, our study also illuminated the therapeutic potential of OB stimulation. OB stimulation noticeably reduced plaque accumulation in Aβ animals not only within the OB but also across other brain regions, including the PFC, dHPC, vHPC, and EC (Aβ+Stim vs. Aβ, p < 0.001). This widespread reduction in plaque accumulation points towards a broader protective effect of OB stimulation, suggesting its ability to modulate Aβ-related neurodegenerative processes beyond the OB itself. These findings collectively emphasize the complex interplay between Aβ, cellular viability, and the potential of OB stimulation as a therapeutic avenue in countering the deleterious effects of Aβ accumulation in multiple brain regions.

### 3.3 OB stimulation augmented the brain regions activity in AD-like animals

Power of LFP signals serves as a reflection of brain activity patterns during cognitive tasks.^34^ Alteration in the power of brain regions oscillations, particularly in higher frequency bands, has been documented in both patients and animal models of AD.^35^ In this analysis, our primary objective was to examine the potential influence of OB stimulation on brain rhythms while engaged in a working memory task. Additionally, we sought to discern potential differences in power patterns within the OB-EC-HPC-mPFC circuit between instances of correct and wrong task performance (Fig. 5).

**Figure 5.**
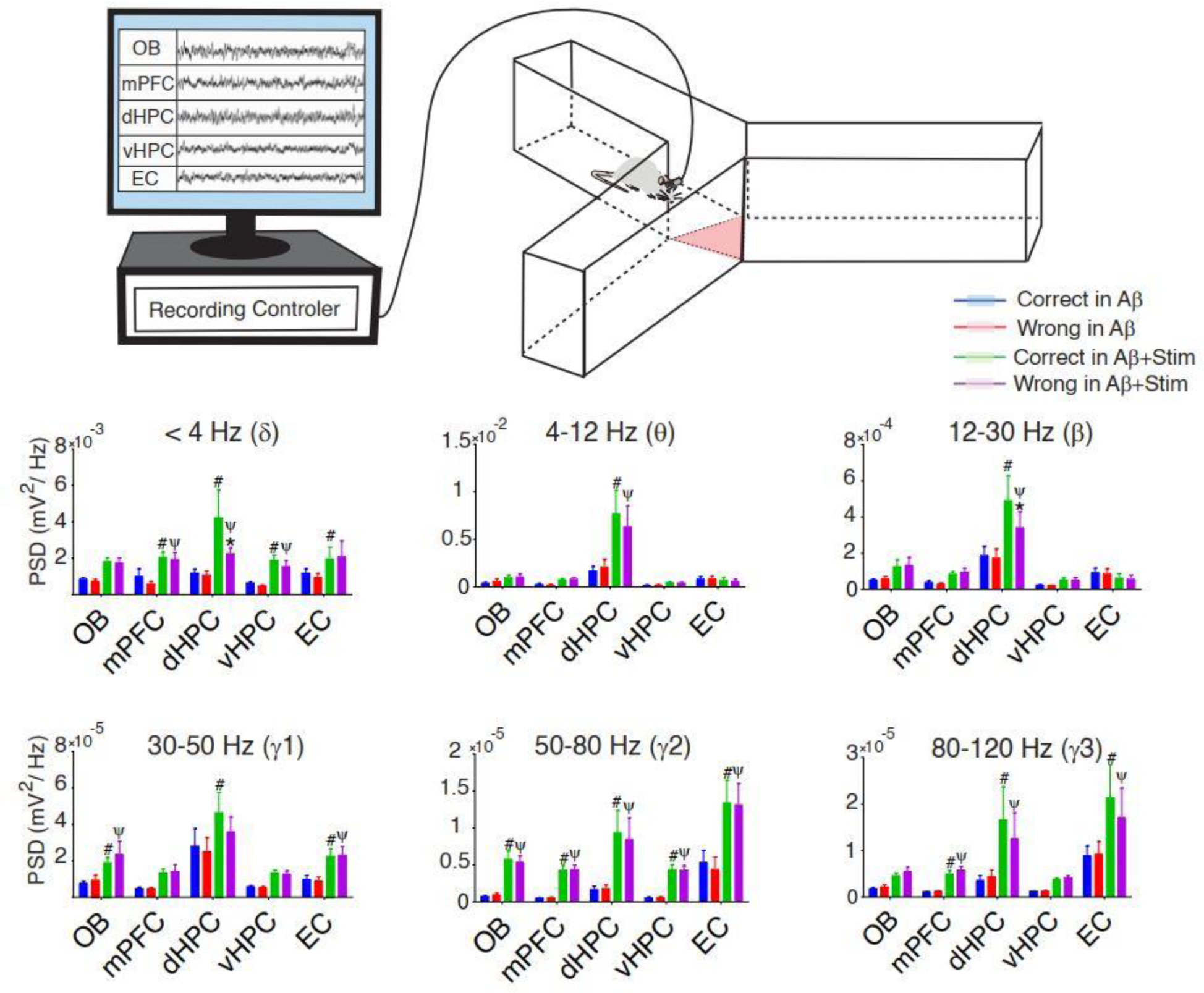
Power spectral density of brain regions during working memory task. (Top) A schematic illustration of the rat while exploring in Y-maze. The red triangle is the central zone in which LFP signals of animals when exploring this area are considered for further analysis. (Bottom) Average PSD during working memory performance. OB stimulation of Aβ animals amplified PSD during working memory decisions compared to Aβ without stimulation. n=6. Data were analyzed by two-way repeated measures ANOVA; * Correct vs Wrong within the group; ψ Correct and # Wrong between groups. PSD, Power spectral Density; OB, olfactory bulb; mPFC, medial prefrontal cortex; dHPC, dorsal hippocampus; vHPC, ventral hippocampus; EC, entorhinal cortex.

Our investigations indicated that OB stimulation significantly amplified power of the mPFC, HPC and EC at the delta band (< 4 Hz) when Aβ animals made the correct decision during working memory task (Aβ+Stim vs. Aβ, p < 0.05). At this frequency band, within-group analysis showed that the power of signals during correct trials was higher than wrong trials in in the dHPC of Aβ animals (p < 0.05).

At theta (4-12 Hz) and beta frequency range (12-30 Hz), OB stimulation of Aβ animals significantly augmented power in dHPC during working memory task (Aβ+Stim vs. Aβ, p < 0.05). At beta frequency, OB stimulation led to higher significant values of power in the dHPC during correct trials compared to the time that Aβ animals were meant to select the wrong arena (p < 0.05).

OB stimulation resulted in an augmentation of power in OB, dHPC and EC at gamma-1 (30-50 Hz), in the all five brain regions at gamma-2 (50-80 Hz), and in mPFC, dHPC and EC at gamma-3 (80-120 Hz) when Aβ animals performed working memory task (Aβ+Stim vs. Aβ, p < 0.05).

Taken together, our results suggest that OB stimulation in Aβ animals can increase the activity of the signals in the OB-mPFC-dHPC-vHPC-EC network.

### 3.4 OB stimulation increased functional connectivity of brain regions in AD-like animals

We have previously reported that functional connectivity in the network between OB, mPFC and HPC is necessary for making the correct decision during working memory performance.^11,12^ Therefore, after addressing the effect of OB stimulation on the activity of the network regions, here we intended to find the strength of coherency between OB, mPFC, vHPC, dHPC and EC due to applying OB stimulation in Aβ animals. As shown in figure 6, the network between the five mentioned brain regions was more strongly connected in the OB stimulated Aβ animals especially in higher frequency bands during working memory task.

**Figure 6.**
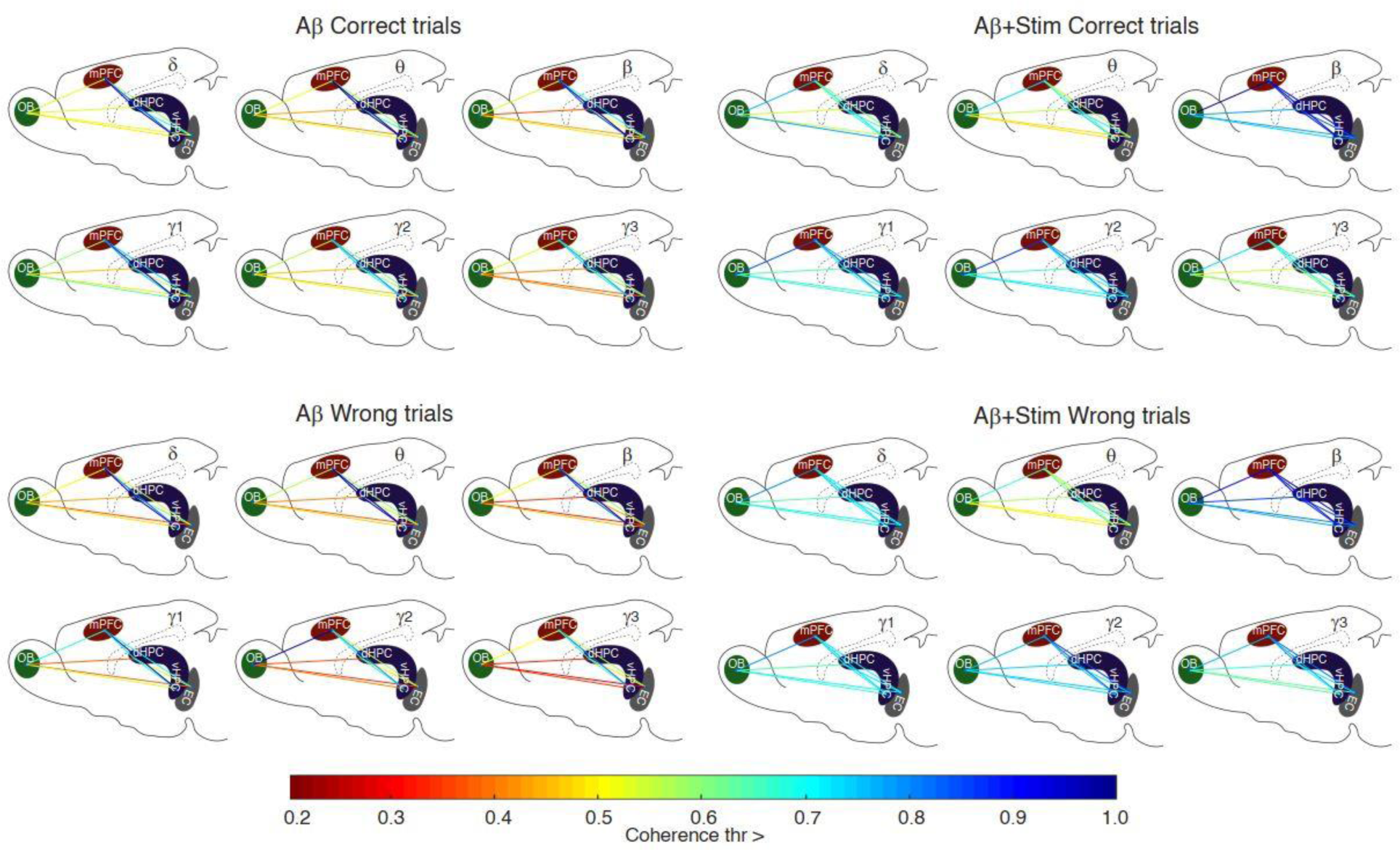
Intensity of functional connectivity in the network between OB, mPFC, vHPC, dHPC and EC during working memory performance. Five brain regions are connected according to the threshold of coherence. Different colors of the lines from dark red to blue indicate the intensity of connection between pairwise regions that has the same or higher value than the designated value for each threshold from 0.2 to 1. Correct and wrong represent the time of the signals that animals were exploring the center of the maze before selecting the correct or wrong arena. OB stimulation increased the strength of coherence within the network during working memory task particularly at higher frequency bands. OB, olfactory bulb; mPFC, medial prefrontal cortex; dHPC, dorsal hippocampus; vHPC, ventral hippocampus; EC, entorhinal cortex.

## 4 Discussion

DBS for treating AD is relatively new.^36^ Although evidence highlights its benefits for memory impairments,^19^ the invasiveness of the surgical procedure and associated side effects hinder extensive research.^37^ Therefore, exploring less invasive approaches to access deep brain areas could be a promising step in treating cognitive diseases like Alzheimer’s. Considering the OB’s connections with memory-associated brain regions and its accessibility via the nasal cavity, we hypothesized it could be an ideal candidate for electrical stimulation to reduce memory deficits induced by AD. Our study demonstrated that electrical stimulation of the OB effectively prevented the development of working memory disorders in AD-like model rats. This was likely due to reduced Aβ accumulation in the brain regions, improved OB cell viability, and enhanced power and functional connectivity within the brain regions involved in working memory networks. To our knowledge, this preclinical study is the first to explore the impact of OB stimulation in alleviating AD progression.

The OB plays a significant role in working memory performance.^11,12^ A significant mechanism through which the OB potentially impacts working memory is nasal breathing.^38,39^ The OB acts as a gateway, allowing nasal respiration to dynamically modulate brain activity.^39^ This specialized role is enabled by the responsiveness of olfactory sensory neurons (OSNs) to both chemical signals and mechanical changes in air pressure during nasal breathing cycles.^40^ These processes lead to respiratory entrained oscillations in the OB and associated areas such as EC, HPC, amygdala, and PFC,^41^ potentially impacting cognitive functions like memory.^41^ Studies indicate that memory tasks yield better results when engaging in nasal breathing, emphasizing the olfactory pathway’s pivotal role in memory processes.^42^ Disrupting this pathway by shifting respiration from the nose to the mouth or inhibiting the OB could eliminate respiratory-induced oscillations, impacting cognitive processes.^42-44^ Additionally, inflammation in the nasal mucosa disrupting OSNs function could reduce hippocampal-prefrontal coupling and impair working memory in rats.^25^ Given that the OB is one of the first brain regions where Aβ accumulation and cell loss occur in the early stages of AD,^45^ it becomes an attractive target for preventing working memory disorders in AD. Our previous study utilizing nasal air-puffing to stimulate the OB in rats showed improved working memory performance.^25^ In the current study, administering electrical stimulation to the OB in conjunction with AD model development resulted in the prevention of working memory deterioration in animals. OB stimulation likely modulated neural dynamics and activity patterns in memory areas and reduced Alzheimer’s-related pathology within memory networks. Through these effects, electrical stimulation of the OB might mitigate declines in working memory associated with AD.

Our results revealed that electrical stimulation of the OB in AD-like model rats enhanced oscillatory activity and functional connectivity within the OB-EC-HPC-mPFC circuit during working memory tasks. Pathophysiological aggregation of Aβ during AD development can manifest as aberrant electrical activity across various brain regions, resulting in a wide range of cognitive impairments.^34^ Disturbances in electrical activity often manifest as reduced oscillations in different frequency bands and diminished synchronization between brain regions.^35^ For instance, alterations in the gamma frequency band in the hippocampus of AD model rats can disrupt spatial working memory task performance.^46^ Reduced synchronization at this frequency is critical for memory performance.^47,48^ Altered power of delta, theta and beta oscillations has also been noted in patients with AD,^35^ disrupting the functional network in frontal and temporal regions and potentially causing working memory deficits.^18,34^ The hippocampal theta power reduction could disrupt cortico-hippocampal connectivity, altering memory performance.^34^

Therefore, therapeutic interventions targeting AD often focus on altering brain oscillation patterns and functional connectivity between brain regions.^49^ Electrical stimulation-based methods have the potential to improve abnormal oscillatory patterns, particularly in regions associated with the hippocampus.^50^ In our previous study, OB stimulation through nasal air-puffing enhanced correlation and coherence between OB and HPC.^26^ Working memory tasks exhibit specific patterns of oscillatory activity and functional connectivity between the HPC, mPFC, and EC.^51,52^ Synchrony between the hippocampus and PFC is associated with successful spatial working memory performance.^51,52^ Accordingly, in our study, OB electrical stimulation improved working memory performance by alleviating the power and functional connectivity of memory circuits. These observations likely stem from the same mechanisms supporting the effectiveness of DBS in modulating brain activity, including increased neurogenesis, alterations in plasticity, and the establishment of new connections across various brain networks.^53-55^ Stimulating the brain can increase cell counts in memory-linked areas, including the hippocampus, by counteracting disturbances in neurogenesis and speeding up cell proliferation.^56^ It may also release neurotransmitters and alter synaptic activity in these regions to enhance memory in AD.^55,57,58^ Our study observed an enhancement of cell viability in the OB stimulation group, suggesting that OB stimulation might improve working memory through these mechanisms. Overall, architectural modifications along with the prevention of Aβ accumulation and enhancement of cell survival in the OB might restore the brain’s electrical activity to a more balanced state.

In Aβ animals, interregional coherence was reduced compared to the stimulation group. OB electrical stimulation appears to have prevented circuit disturbances, resulting in improved working memory performance in the stimulation group. A prior study involving healthy animals demonstrated that coherency between the OB and the PFC-HPC circuit increased during correct task performance.^11^ This indicates that OB electrical stimulation brought the working memory circuit’s activity closer to that of healthy animals. This impact of olfactory pathway stimulation was also evident in comatose patients.^59^ Olfactory pathway stimulation at regular respiration frequency led to increased activity of gamma oscillations (30–100 Hz) and functional connectivity in the default mode network.^59^ Gamma oscillations are crucial for integrating and combining information within local circuits and across different brain regions in healthy individuals.^60,61^ Therefore, olfactory pathway stimulation seems to alleviate cognitive functions by modulating brain activity, especially in gamma ranges.

Another mechanism through which electrical stimulation of the OB could prevent working memory disorders is by mitigating the deposition of Aβ plaques in the OB-EC-HPC-mPFC network. Aβ deposition is directly related to the reduction of working memory capacity.^62,63^ Alleviating Aβ deposition in the OB is particularly important because the OB is a hub for adult neurogenesis.^64^ The process of adult neurogenesis in the OB involves specialized neural stem cells that give rise to neural precursor cells, eventually differentiating into mature neurons.^65^ These newly generated neurons not only integrate into existing neural circuits within the OB but also contribute to the brain networks engaged in cognitive functions like working memory.^43,66^ On the other hand, Aβ proteins exhibit prion-like characteristics.^67^ After aggregating in the OB, these proteins can spread to other connected regions, including the EC and HPC, impairing working memory mediated by these downstream areas.^67^ In our study, electrical stimulation of the OB in Aβ animals enhanced surviving cells in this area and concomitantly prevented Aβ deposition. It may be concluded that OB stimulation, by intensifying neurogenesis, preventing cell loss, and ameliorating Aβ accumulation, improved working memory in AD model animals.

Therefore, OB stimulation might impact working memory via preventing plaque buildup and increasing neurogenesis locally in the OB, as well as reducing plaque accumulation in the interconnected OB-EC-HPC-mPFC circuit. Furthermore, OB stimulation could potentially safeguard working memory by mitigating inflammation and apoptosis within the working memory network.^26^ Although this study didn’t investigate these complexities, a prior study conducted by us demonstrated that OB stimulation, achieved through air puffing, effectively prevented hippocampal inflammation and apoptosis in mechanically ventilated animals.^26^

This study didn’t explore the molecular and cellular mechanisms underlying the alteration in brain electrical activity resulting from OB stimulation. Further investigations are warranted to elucidate the precise microscopic processes through which olfactory pathway stimulation modulates neuronal population activity. In our investigation, OB electrical stimulation led to a decrease in Aβ aggregation in AD model animals. However, it should be mentioned that the stimulation did not completely revert Aβ levels to those observed in control animals. This limitation might be linked to the relatively short duration of the stimulation sessions (1 hour daily) in our experimental design. Conceivably, implementing continuous electrical stimulation throughout the entire day, akin to the methodology employed in clinical DBS procedures,^19^ may yield greater effectiveness in preventing Aβ deposition. Additionally, prospective alterations in olfactory perception were not examined in this study. Given the potential impact of altered smell perception on cognitive functions, including memory,^68^ forthcoming studies employing olfactory pathway stimulation should incorporate behavioral evaluations to ensure the olfactory system’s functionality within the research design. However, in a human study, electrical stimulation of OSNs under threshold value altered the activity of olfactory-related brain regions, whereas participants did not perceive any sense of smell.^69^ The study concluded that indiscriminate stimulation of the olfactory epithelium is insufficient to trigger a sense of smell.^69^

## 5 CONCLUSION

This preclinical study focused on early stages of AD and demonstrated the effectiveness of high-frequency electrical stimulation of the OB in preventing AD-related pathological, behavioral, and electrophysiological issues. Particularly significant is the potential of olfactory pathway stimulation for modulating deep brain activity, emphasizing its relevance in developing preventive strategies for AD, especially targeting individuals in the MCI stage. Given the accessible nature of the olfactory pathway through the nasal cavity, this study opens new possibilities for less invasive brain stimulation methods. Electrical stimulation of OSNs via the nose holds promise as an effective approach, potentially revolutionizing treatments for neurocognitive dysfunctions like AD, depression and epilepsy.

## AUTHOR CONTRIBUTIONS

Conceptualization: Morteza Salimi, Mohammad Reza Raoufy, Mohammad Javan and Javad MirnajafiZadeh. Methodology, Validation: Morteza Salimi, Mohammad Reza Raoufy and Milad Nazari. Investigation and data collection and analysis: Morteza Salimi, Mohammad Reza Raoufy, Milad Nazari, Samaneh Dehghan. Investigation and data collection and analysis: Morteza Salimi and Milad Nazari. Writing–original manuscript: Morteza Salimi, Mohammad Reza Raoufy, Milad Nazari and Payam Shahsavar. Manuscript review and editing: Morteza Salimi, Mohammad Reza Raoufy and Payam Shahsavar

## ACKNOWLEDGMENTS

We extend our gratitude to Mr. Reza Vaziri, Mohsen Sharifi, Farhad Farkhondeh, and Forough Foolad for their valuable technical support in this article. Furthermore, we would like to acknowledge and appreciate the exceptional assistance provided by Mr. Alireza Mani, for which we are profoundly thankful.

## CONFLICT OF INTEREST STATEMENT

All authors report no conflict of interests.

## DATA AVAILABILITY STATEMENT

The corresponding author can provide the data supporting the study findings upon a reasonable request.

## FUNDING

This work was supported by Tarbiat Modares University (grant number IG-39709).

## ORCID

Mohammad Reza Raoufy* https://orcid.org/0000-0002-9292-8564

## Notes

### Competing Interest Statement

The authors have declared no competing interest.

